# Inducible *in vivo* genome editing in the sea star *Patiria miniata*

**DOI:** 10.1101/2023.01.09.523328

**Authors:** Olga Zueva, Veronica F. Hinman

## Abstract

For centuries, echinoderms, a marine-invertebrate phylum, have fascinated scientists for their developmental and postembryonic phenomen. Experimentation on their eggs and embryos in particular have contributed foundation scientific advances. However, powerful molecular genetic studies are restricted to embryonic developmental stages which are amenable to genetic perturbation by microinjection of reagents into the zygotes. This represents a significant bottleneck to the study of postembryonic processes in where the earliest function of a gene must remain intact. We therefore sought to establish a spatio-temporal turnable gene editing tool for these species. Here, using the sea star Patiria miniata as a model we introduce a chemically inducible, Tet-ON, gene expression system. Pairing this Tet-ON system with CRISPR-mediated gene alteration technology we show as a proof-of-principle demonstration an inducible gene editing in the sea star transgenic cell populations for the first time in echinoderm biology. The approach we show here can be adapted for use in other species of echinoderms and will also extend experimental possibilities tremendously.

## Introduction

The extraordinary diversity of physiologies, behaviors and morphologies across the metazoan tree of life presents an opportunity to explore and understand fundamental mechanisms underpinning an array of biological phenomena. Krogh, 1929 famously stated that for any biological process there will be some species most suited for its study. This principle underlies model organism research and has driven biological experimentation and especially biomedical advances. Echinoderms such as sea urchins and sea stars, and in particular their early embryos and larvae, have been model organisms for centuries (Ernst, 1997). Echinoderm research has contributed to our understanding of foundational concepts in biology such as, chromosomal inheritance (Laubichler and Davidson, 2008), phagocytosis (Rather, 1970), induction and regulative development (Angerer and Angerer, 1999), and gene regulatory networks function and evolution (Cary et al., 2020; Davidson et al., 2002; Hatleberg and Hinman, 2021; Hinman and Cheatle Jarvela, 2014; Hinman and Davidson, 2007; Hinman et al., 2009).

Echinoderms, however, also exhibit a wealth of fascinating postembryonic phenomena. For example, many species of echinoderms have extraordinary capacities for regeneration. Brittle stars are well known for arm regeneration upon amputation (Czarkwiani et al., 2022; Mashanov et al., 2022), some sea star species can regrow complete organism from remnant arm (known as comet sea star), some sea cucumber species are able to regenerate full organism after fusion and from the remnant body part (Dolmatov, 2021). Echinoderm larvae also have outstanding capacity to undergo whole body regeneration (Wolff and Hinman, 2021). Additionally, many echinoderms also undergo dramatic metamorphosis, transitioning from a free swimming planktonic larvae, to a benthic juvenile following extensive, often with very rapid morphogenesis. Some species of echinoderms are extremophiles living in antarctic seas and in deep ocean vents (Chen et al., 2022; Zhang et al., 2022). While fascinating, very little is known about genetic control of these processes because there are not yet tools to genetically edit post embryonic stages whilst preserving early developmental processes.

The ability to perturb gene function at a specific time and in a specific cell type is vital for understanding biological mechanisms. Overexpression, chemical inhibition, reporter protein expression, knockdown via antisense morpholinos, RNAi and knockout of genes of interest have all been used in echinoderms functional studies (Buckley and Ettensohn, 2019; Lin et al., 2019; Molina et al., 2019). Three major genome editing technologies ZFNs, TALENs, and CRISPR/Cas have been applied to target sea urchin genes, but CRISPR is the most versalite, low-cost, easily multiplexed tool for echinoderm functional genomic studies.

CRISPR/Cas - mediated gene editing provides a wide variety of tools for genetic perturbation. Cas9 endonuclease is directed by an RNA guide to cut a target DNA sequence to enable facile editing of the genome in almost all organisms. This can be applied to non coding regulatory or coding regions of the genome, with Cas9 and an RNA guide in a cell, the ribonucleoprotein complex (RNPs) will cleave its DNA target, irrespective of the type, environment, internal state or developmental stage of the cell. The editing strategy relies on complex DNA repair pathways and DNA double-strand breaks. In the simplest application of the technology, the gRNA directed Cas9 targets the genome and causes mutagenesis by cleavage and imperfect repair to result in an edited gene locus. CRISPR/Cas - based genome editing enables the rapid genetic manipulation of any genomic locus and this technology opens up the opportunity to study not only gene function but also the roles of non coding DNA. However, an additional challenge remains that many genes of interest, and especially regulatory genes, are highly pleomorphic and studying later, post embryonic process requires a fine spatio-temporal control of gene editing so that earlier and other non relevant processes are not functionally changed. Pairing CRISPR/Cas gene editing with an inducible system is one way to provide spatial, and temporal control to gene editing. We therefore sought to establish a spatio-temporally inducible CRISPR/*Sp*Cas9 gene editing system to study postembryonic processes in echinoderms, using the sea star, *Patiria miniata* as our model system.

CRISPR/Cas9 - based genome editing was recently successfully applied in echinoderm (Lin et al., 2019; Oulhen et al., 2022), which represents a major step towards more sophisticated genetic manipulation, and the establishment of reproducible transgenic adult lines. Before CRISPR technological era the genetic perturbation studies in echinoderm biology heavily rely on using MASO (morpholino antisense oligonucleotides) knockdowns, which are not long lasting and therefore do not perturb gene expression during post embryonic and adult stages. Here we demonstrate inducible *in vivo* genome editing with CRISPR/*Sp*Cas9 paired with the chemically inducible Tet-ON/OFF system. This system has been developed over 25 years ago from components of the bacterial transposon Tn10-encoded Tet operon (Hillen and Berens, 1994; Wissmann et al., 1986). Tet-regulatable systems are based on the Tet repressor protein (TetR) and the Tet operator DNA sequence (TetO DNA). In the absence of tetracycline or its derivative doxycycline (DOX), the TetR protein binds to the TetO DNA sequence. When tetracycline is present TetR changes its conformation and detaches from the DNA. Tet-ON and Tet-OFF regulatory systems have been developed to induce or repress gene expression and fusion of TetR to the VP16 transcriptional activator of herpes simplex virus allows for control in eukaryotes (Kang et al., 2019). In this study we used the commercially available third generation Tet-ON (Clontech) system. This system is based on two components: Tet-ON 3G, a reverse transactivator protein (rtTA) that binds the TRE3Gs DNA sequences, the Tetracycline Responsive Elements (TREs) only in the presence of DOX (Figure 1A). Positioning *Sp*Cas9 endonuclease expression under Tet-ON control, proof - of - concept experiments demonstrated the effectiveness of an inducible CRISPR/Cas9 gene editing system in the sea star *P. miniata* (Figure 2).

**Fig. 1.**
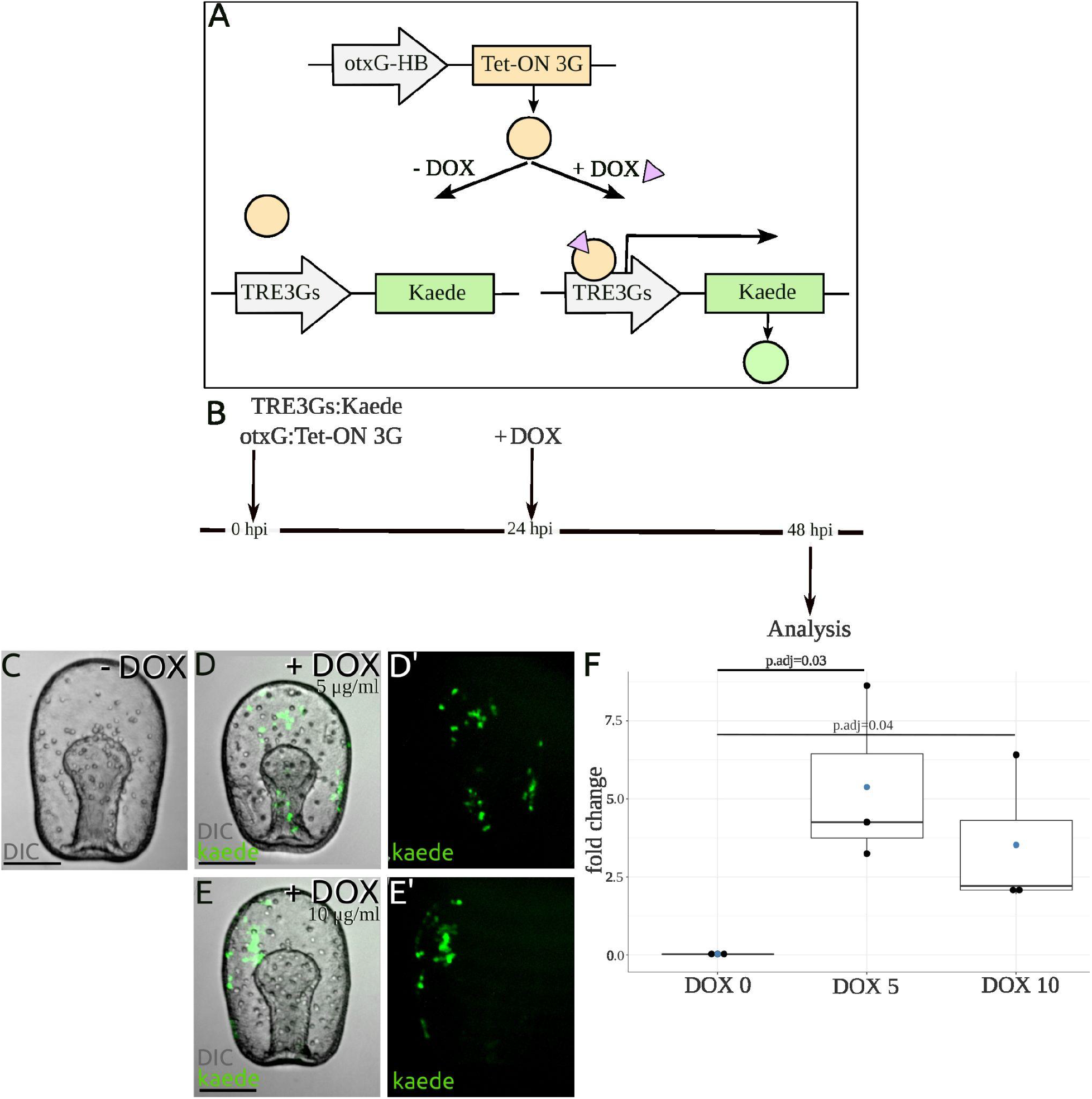
Plasmid based Tet-ON inducible expression system in the sea star *Patiria miniata*. (A) Schematic of doxycycline (DOX) - regulated expression system. In the absence of DOX, the modified reverse Tet repressor (Tet-ON 3G) is unable to bind to the Tet operator (TRE3Gs). Upon addition, DOX binds to the Tet-ON 3G rendering it able to bind to the TRE3Gs. As a result, the gene of interest is transcribed. Tet-ON 3G is a fusion protein made from the modified reverse Tet repressor and three repeats of viral transcriptional activator (VP16), which will activate transcription from a minCMV promoter with a seven repeats of Tet operator (TRE3Gs) (Table S1). *Tet-ON 3G* gene expression is under *Pm*-specific enhancer (otxG) and basal RNA polymerase II promoter (HB). (B) Schematic of experimental design. DOX was added at sea water after 24 h post injection (hpi). (C-F) Evaluation of the inducible expression system (A) in *P. miniata* embryos (Table S2). Z - projections of confocal stacks of in vivo images. Scale bars: 50 μm. (C - E’) Kaede protein expression in the absence (C) or in the presence of DOX at 5 μg/ml (D-D’) and 10 μg/ml (E-E’). (F) Relative quantification of *kaede* gene expression in response to two concentrations of DOX at 5 μg/ml (DOX5) and 10 μg/ml (DOX10). Three RT-qPCR reactions (Table S3) are carried out for each sample, normalized against *laminin2b receptor*. The significance of fold difference upon DOX induction is calculated by Student’s *t*-test (*n* = 3 biological replicates, ^∗^*p* < 0.05). Boxplots showing individual data points (black), mean values (blue) and pairwise P values.

**Fig. 2.**
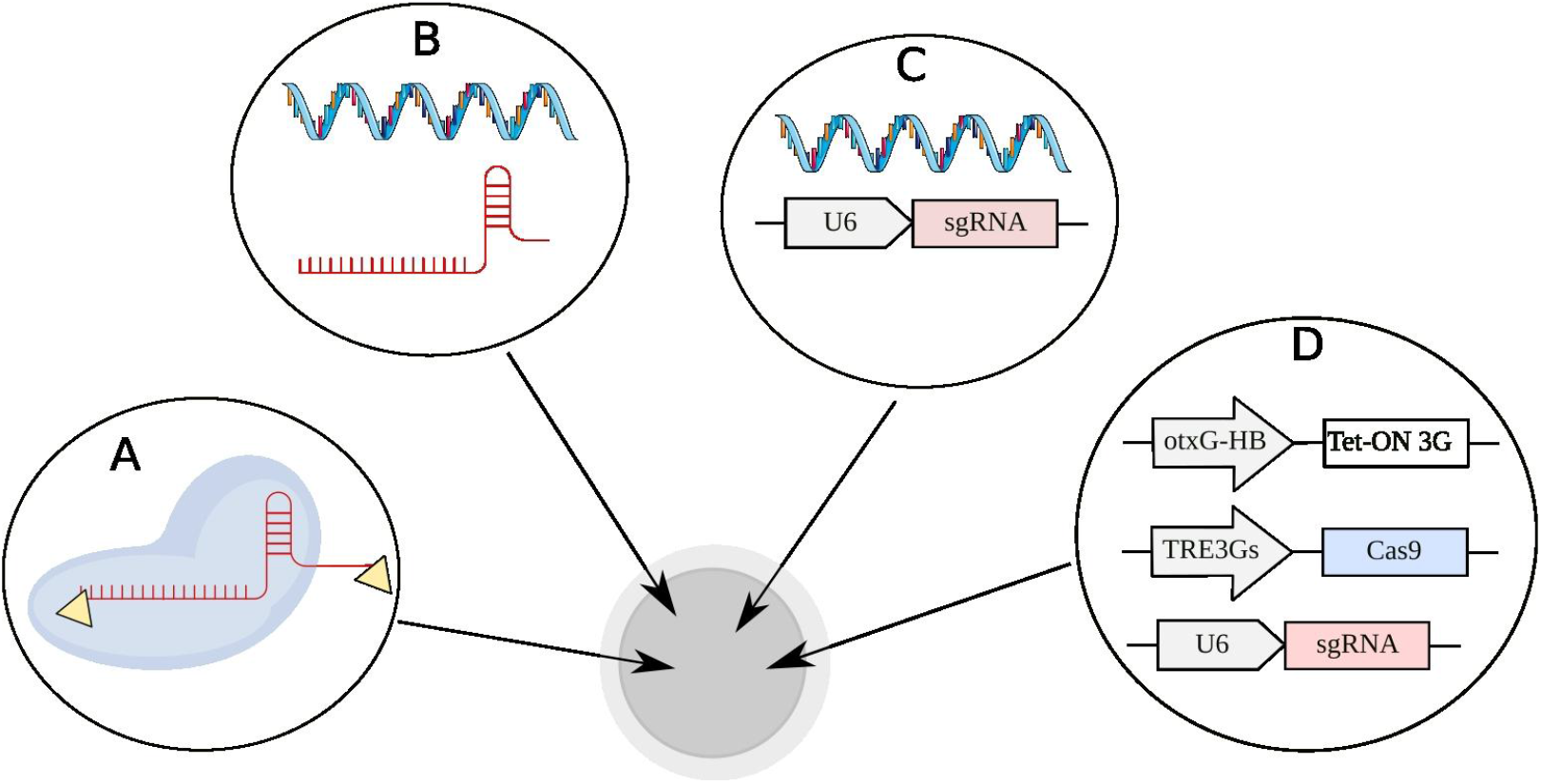
Four strategies of CRISPR delivery into the fertilized egg of *P. miniata*. (A) Synthego *Sp*Cas9 nuclease complexed with synthetic sgRNAs (Synthego) with 5’ and 3’ ends chemical modifications. (B) Human-codon optimized *Sp*Cas9 (hCas9) mRNA with unmodified, in vitro transcribed sgRNA. (C) hCas9 mRNA with plasmid - expressed sgRNA under *Pm*-specific U6 regulatory elements. (D) Spatio-temporal control system for genome editing using induced CRISPR (iCRISPR) in otxG positive cells. Some illustrations were created with bioicons.com.

Four different strategies for introducing the CRISPR system into the sea star cells were tested, the editing efficiency of *Delta* (*Dll*) locus was validated and phenotypic outcomes upon mutagenesis were assayed. The effectiveness of inducible CRISPR (iCRISPR) knockout was compared with the most effective in vitro complexed RNPs strategy of introducing the system into the cells. In order to express the guide RNAs, which are RNA polymerase III transcribed, we determined the *P. miniata* regulatory elements of U6 small nuclear RNA. This is the first example of functional *in vivo* application of Pol III - driven small RNAs expression in echinoderm biology.

Our iCRISPR system functions in a clonal mosaic fashion, wherever the Tet-ON 3G (rtTA) expresses and thus the Cas9, which allows to specify genes manipulation to defined tissue/cell populations. Using U6-driven sgRNAs targeting several genes would allow simultaneous manipulation of different genes in a given tissue/cells. We think that this inducible two component plasmid system has great potential to transform *Patria miniata*, and other species of echinoderms, for exploration of the genetic underpinnings and suite of postembryonic developmental processes.

## Results

### Construction of the Tet-ON reporter system and efficiency validation

Two plasmid-based vectors were built to test the Tet-ON system in *Patiria miniata* (Figure 1A). One vector carries the Tet-ON 3G transactivator and another the fluorescent reporter Kaede. The expression of the transactivator (Tet-ON 3G) was put under control of the *P. miniata* regulatory element (*Pm*otxG) upstream of the minimal *Strongylocentrotus purpuratus* hatching enzyme RNA polymerase II promoter, HB (Wei et al., 1995). The otxG enhancer was previously identified as the cis-regulatory element responsible for control of *Am*otxβ1/2 expression during blastula and early gastrula stage in the ectodermal and endomesodermal cells that drives expression broadly throughput the ectoderm and endoderm by hatching stage (Figure S1, (Hinman et al., 2007)). Tet-ON 3G is a fusion protein of third generation rtTA and three viral VP16-derived minimal activation domains (Table S1). TRE3Gs contains seven TetO sequence repeats with a 36 bp center-to-center distance upstream of a minimal CMV promoter (Table S1). Upstream of a fluorescent reporter (Kaede) are TRE3Gs. Both circular plasmids were delivered into the fertilized eggs by microinjection at the final concentration of 10 ng/μl at 1:1 ratio in 120mM KCl injection solution.

Expression of Kaede protein should only be detected when effective doses of DOX are applied to the two vector system (Figure 1A). To determine the optimal DOX concentration for activation of reporter expression in the sea star (Figure 1B) two commonly used concentrations of drug/inducer were tested, 5 and 10 μg/ml (Campbell et al., 2012). DOX of 5μg/ml (DOX5) and 10 μg/ml (DOX10) was added to the seawater 24 hours post injection (blastula stage). The control group (DOX0) received only an ethanol carrier (1/1000 dilution). 24 hours after addition of DOX (mid gastrula stage), 300 embryos per dish in three independent biological replicates of DOX0, DOX5 and DOX10 were assayed.

The expression of Kaede was assessed by fluorescence microscopy (Figure 1C-E’; Table S2) and qPCR (Figure 1F; Table S3). In the presence of DOX5 green fluorescence was seen in 121.7 ± 33.3 embryos and in 92.3 ± 20.4 embryos upon DOX10 addition in three independent biological replicates (Table S2). Fluorescence was detected in multiple clonal patches in the ectoderm and endoderm. Plasmid constructs are thought to be stably inherited after a few cell divisions and thus express mosaically in clones of cells. The observed Kaede localization reflects the fluorescent pattern driven by otxG enhancer directly regulating reporter with the same basal promoter (Figure S1). In the presence of DOX, Kaede mRNA was expressed at 5.4 ± 2.9 (DOX5), 3.5 ± 2.5 (DOX10) - fold higher levels than in the case without antibiotic induction (Figure 1F; Table S3).

A series of control experiments were performed to test TRE3Gs leakiness, and to test whether there was any otxG enhancer cross-reactivity with Kaede in the two vector system (Table S2). TRE3Gs leakiness test was conducted by injection TRE3Gs:Kaede only. No Kaede positive embryos were observed during this experiment. It is thought that injected plasmid DNA is concatenated during integration into the genome; it is possible that reporter expression might be driven by another vector’s regulatory element. To test otxG enhancer cross-reactivity we amplify otxG:TetR region without VP16 domains (Table S1) and inject this linear DNA amplicon with TRE3Gs:Kaede as a circular plasmid followed by DOX5 induction. Only one sick embryo per two independent biological replicates with weak reporter expression was detected during this experiment (Table S2).

In combination these results indicate the Tet-ON system allowed a precise and convenient regulation of gene expression in the sea star.

### Gene editing using CRISPR/Cas9

To establish gene editing in *P. miniata* we first tested three different methods for introducing the Cas9 and sgRNAs into the sea star: (1) pre-assembled ribonuclear complexes (RNPs), (2) Cas9 mRNA with in vitro transcribed sgRNA, and (3) Cas9 mRNA with plasmid-driving sgRNA expression (Figure 2). These methods were evaluated using *Dll* knockout phenotype and sequence analysis. The iCRISPR system was then tested using (1) cell specific, DOX inducible Tet-ON 3G and therefore Cas9 and Kaede reporter and (2) constitutively expressed sgRNAs, to demonstrate spatiotemporal control of gene editing in *Patiria miniata*.

#### *Dll* knockout assay

As a target for knockout, we wanted a non-lethal early developmental, well-characterized gene with a clear phenotype upon manipulation. Delta-Notch signaling pathway which regulates the spatial pattern timing and out-come of cell-fate decisions became our choice. In the sea star Delta-Notch signaling works via lateral inhibition regulatory interactions (Cary et al., 2020). Notch ligand *Delta* gene (*Dll*) is a zygotic gene whose expression is initialized by the 16h early blastula stage reaching the peak of expression by the 24h blastula (McCauley et al., 2015). *Dll* has several roles during development, but most notably loss of *Dll* function leads to an increase of *gcm+* (*glial cells missing*) cells in the ectoderm. In normal embryos *gcm* is expressed in a salt and pepper like fashion with one *gcm*+ cell surrounded by two non *gcm*+ cells. In a *Delta* MASO morphant *gcm* is expressed in adjacent cells (Figure S2). Knockdown or knockout of *Dll* in the sea star also results in excess mesenchymal cells and failure to form coelomic pouches (Cary et al., 2020; Perillo et al., 2022). A *Dll* gene was annotated in the *P. miniata* genome based on orthology from the other species (Foley et al., 2021). The gene consists of 10 exons spanning over 12, 804 bp, has two splice variants and encodes a delta protein ligand that is characterized by a DSL domain, EGF repeats, and a transmembrane domain (Figure 3A).

**Fig. 3.**
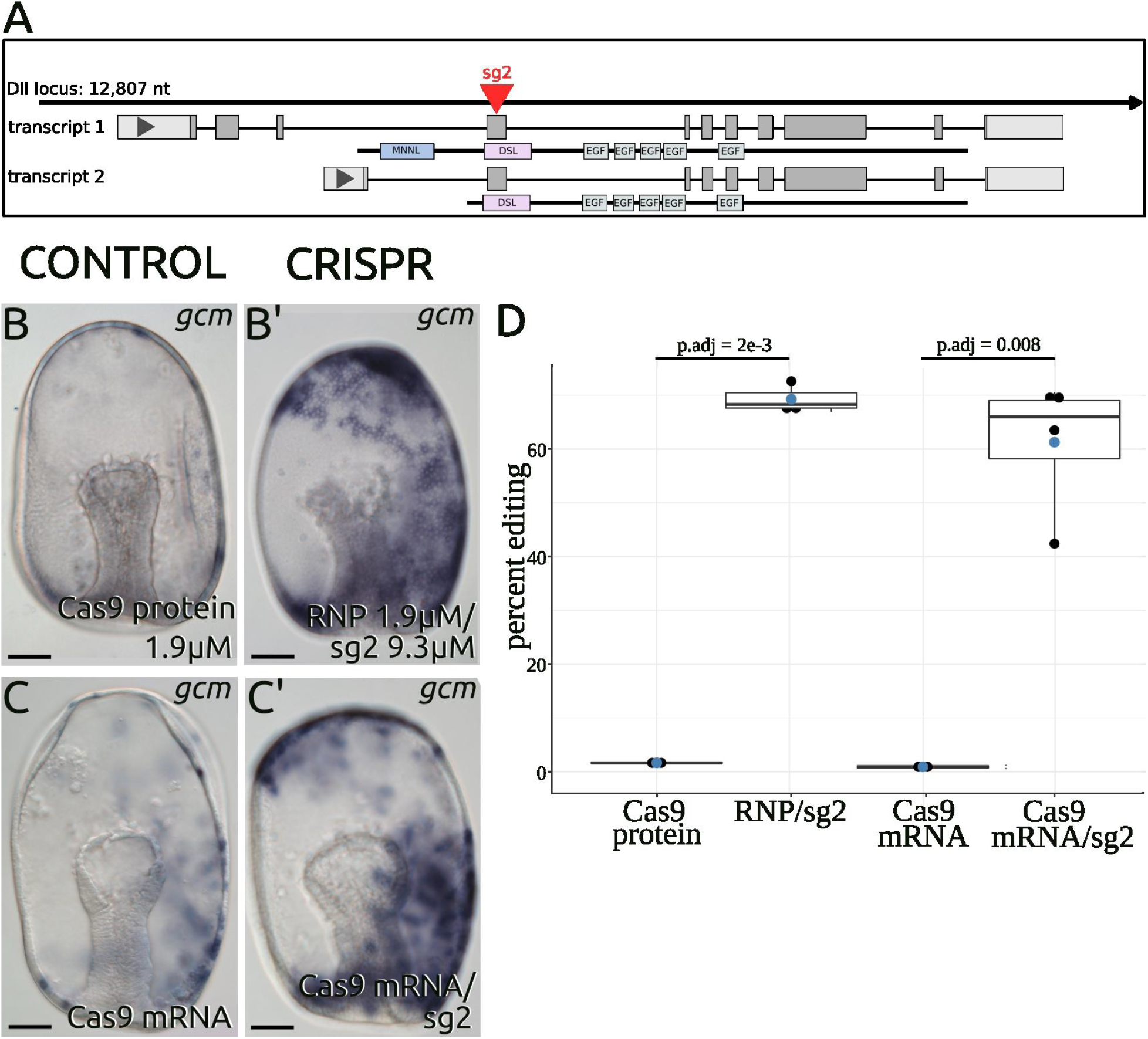
Delta (*Dll*) knockout by CRISPR/Cas 9 technology. (A) Schematic showing the *Dll* locus organization and Cas9 targeting mediated genome editing (sg2). (B - B’) *Sp*Cas9 (wild type with two nuclear localization sites, Synthego) was delivered into the zygotes as a ribonucleoprotein (RNP), complexed with the synthetic sgRNA with chemical modifications (Synthego) in 1:5 the molar ratio. Phenotypic analysis of control (B) and *Dll*-KO (B’) embryos after the delivery of RNP visualized by WMISH against *gcm. Dll*-KO embryos showing strong overexpression of *gcm*. This effect was shown previously by using *Dll* MASO perturbation (Figure S2). (C - C’) In order to generate an inducible CRISPR construct we tested the efficiency of human codon optimized *Sp*Cas9-2xNLS (addgene # 72602 with SV40 polyA) mRNA coinjected with unmodified, in vitro transcribed *Dll* sgRNA (sg2). WMISH against *gcm* performed on control (C) and *Dll*-KO (C’) embryos. (D) The quantified percent editing in CRISPR embryos (RNP/sg2, and Cas9 mRNA/sg2) was statistically significant relative to unedited embryos (Cas9 protein, and Cas9 mRNA) with the confidence level of 95%. The significance of perturbation is calculated by Student’s *t*-test. Mutation efficiency is assessed by single embryo genotypic analysis. At least five embryos were genotyped for each experiment, and at least two independent biological repeats were performed. The Indel (insertion - deletions) frequencies and the overall efficiency were calculated using TIDE software (Brinkman et al., 2014). Boxplots showing individual data points (black), mean values (blue) and pairwise P values. Scale bars: 50 μm.

#### *Dll* knockout using synthetic sgRNAs and Cas9 nuclease

Single chimeric gRNAs (sgRNAs) were designed using CHOPCHOP v3 (Labun et al., 2019) to exon 1 (sg1), exon 4 (sg2; shared by both splice variants) (Figure S3 A; Table S1) and synthesized chemically with protecting groups (Synthego). These were mixed with recombinant *Streptococcus pyogenes* Cas9 protein (Synthego) to form ribonuclear complexes (RNPs) in 5:1 molar ratio and delivered into a fertilized egg by microinjection (Figure 2A; 3 B-B’, D; S3). Three different concentrations of RNP complexes (1.2μM Cas/ 6μM sg; 1.9μM Cas/ 9.3μM sg; 3.75μM Cas/ 18.6μM sg) were tested (Figure S3; Table S4). Crispant embryo phenotypes were assayed by colorimetric whole mount in situ hybridization against *gcm* by the 48h gastrula stage (Figure 3 B-B’; S3 B-F). The genomic disruption (indels) at the expected genomic *Dll* locus were assayed by PCR genotyping with primers surrounding the target site at single embryo resolution. Mutation efficiency of both edited and unedited (Cas9 nuclease only) embryos was calculated using Tracking of Indels by DEcomposition (TIDE) software (Brinkman et al., 2014). Reasoning that embryos without RNPs exposure exhibit background levels of indels due to PCR, sequencing errors, genomic polymorphism, the presence of paralogous genes, all control sequences (nuclease only) were subjected to TIDE analysis. In preliminary experiments using RNPs at different concentrations with both sgRNAs (sg1, sg2) we observed concentration dependent editing effects both phenotypically (Figure S3 B-F) and genotypically (Figure S3 G; Table S4). Microinjections of sgRNAs only showed no abnormal phenotype (data not shown). This pilot experiment allows us to choose

RNP at a concentration of 1.9μM Cas9 nuclease with 9.3μM sg2 for following statistically quantified indel frequencies assay (Figure 3D; S4 A; Table S4). In this case we show that the delivery CRISPR system as a RNP complex (Figure 2A) resulted in 68.3 ± 26.5 editing efficiency at *Dll* loci (Figure 3 B-B’, D; S4 A) with greater *gcm* overexpression in ectodermal cells. A 100% editing is not expected due to the heterogenous editing that is likely to occur. Overall, this approach allows us to determine working sgRNA sequences and establish phenotyping, and genotyping assays as the control background.

#### Dll knockout approach using in vitro transcribed sgRNAs and Cas9 mRNA

We next used transcribed sgRNA and Cas9 transcribed mRNA to assess for phenotype and perturbation efficiency (Figure 2B; 3 C-C’, D). We used an optimized tracrRNA scaffold to build chimeric sgRNAs (Table S4), which were transcribed in vitro using the MegaShortScript T7 transcription kit. It has been shown that optimized scaffold with extending the duplex by 4-6 bp and mutation of fourth T to C or G (removing termination signal RNA polymerase III) significantly increase knockout efficiency (Dang et al., 2015); Table S4). Based on the similar codon usage in humans and sea stars, Cas9 sequence codon optimized for human cells was chosen to express the nuclease in the sea star cells. Along with two nuclear localization sequences (NLS) hCas9 mRNA (annotated sequence information in Table S1) was transcribed in vitro, mixed with our in vitro transcribed-sgRNA (sg2) and injected into fertilized eggs. Injected embryos were assayed as from the previous approach. Phenotypic changes visualized by *gcm* in situ hybridization were also observed (Figure 3 C-C’), and editing efficiency judged by TIDE was nearly the same (Figure 3D; Table S4) the first experimental design with RNPs. In case of RNPs the mutation efficiency is 68.3 ± 26.5 % while the transcribed Cas9 and guides produced 61 ± 13 % editing. The slightly diminished effect of the transcribed RNAs may be due to the delayed effect of Cas9 mRNA, and/or instability of home-made sgRNAs. Moreover, activity of different DNA repair pathways likely contributes to the dynamics of indels profiles generated by different Cas9/sgRNA delivery methods (Kosicki et al., 2017). But this assumption remains an unexplored field in echinoderm biology.

#### Constitutive expression of sgRNA under the *Pm*U6

We next needed a system to constitutively express the guide RNAs (Figure 2C). For gRNAs expression we choose to use regulatory sequences from small nuclear RNA U6 genes, which are Pol III - transcribed and have been successfully used to express small RNAs such as siRNAs/shRNAs (Sandy et al., 2005) and gRNAs for CRISPR (Cong et al., 2013) in a broad range of species. The well-characterized human (Hs U6, X07425.1) and sea urchin (Sp U6.1, X76389.1) U6 were used as the queries to search for homologs in the *P. miniata* genome. Five, 87-97% identical matches were found. Reasoning that the cis-elements necessary for transcription of U6 snRNAs are present in the 5’-flanking regions we wanted to identify those regulatory elements. Multiple sequence alignment (Figure 4A) allows us to identify transcription initiation position (+1, with G as the preferred initiation nucleotide), conserved an intragenic control region, and termination signals. However, comparison of the putative regulatory elements showed that the nucleotide sequence diverges significantly. Compared to the location of the TATA box, proximal and distal sequence elements of human and sea urchin sequence to the sea star sequences led us to predict the genomic regulatory region upstream of the *Pm*U6 snRNA coding region. Then to narrow down the potential *Pm*U6 candidate we used open chromatin regions data (ATAC-seq) to identify chromatin accessibility of actively transcribed U6 snRNA at early developmental and regeneration stages. LOC109742214 exhibits open chromatin peak at preference time window (at blastula stage, and at three days post bisection larvae) and was subjected for the following analysis.

**Fig. 4.**
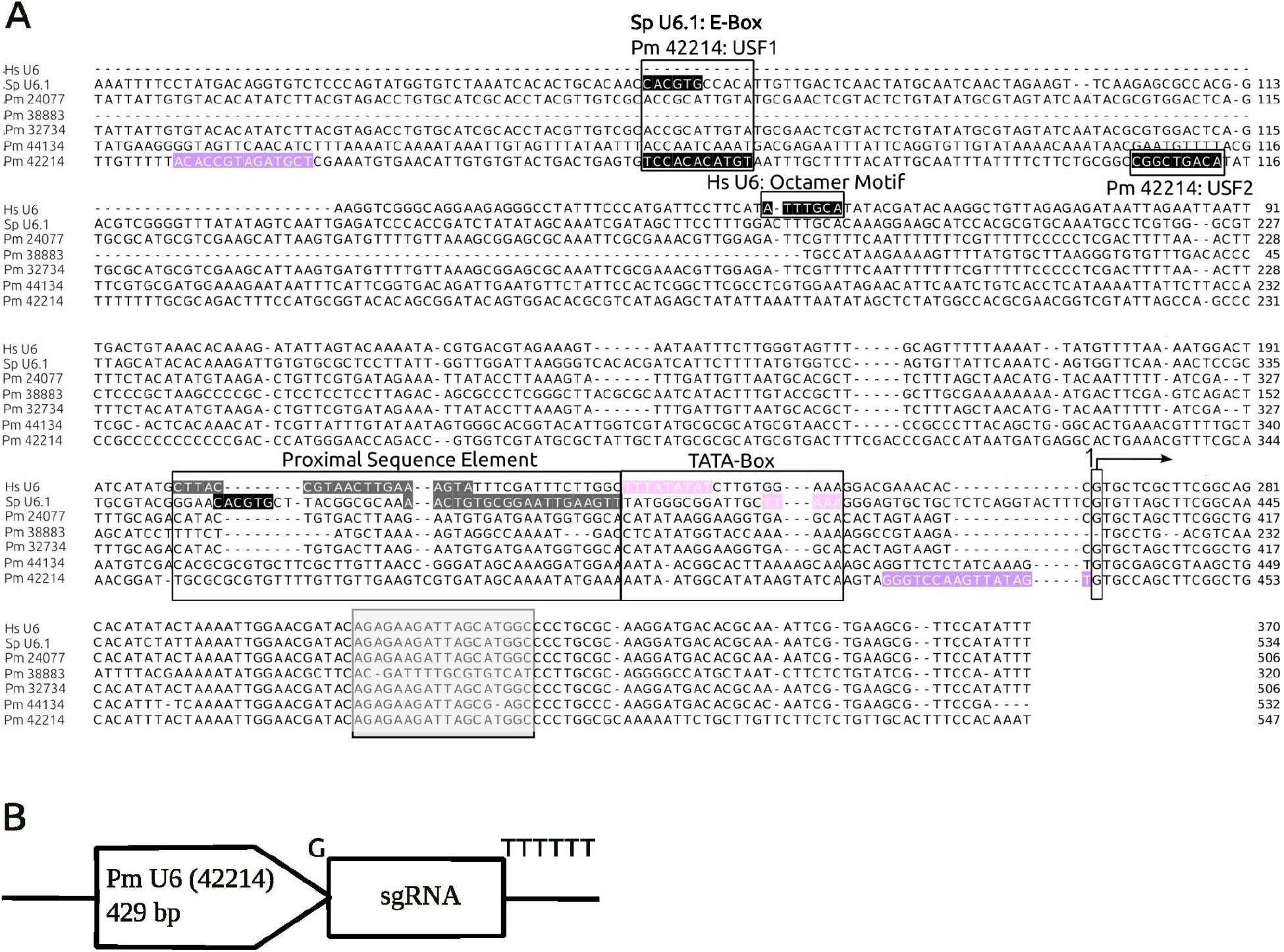
Construction of the *Pm*-specific U6:sgRNA vector. (A) Alignment of the human (Hs U6, X07425.1), sea urchin (Sp U6.1, X76389.1), and five *P. miniata* (LOC119724077, LOC119738883, LOC119732734, LOC119744134, LOC119742214) U6 snRNA genes. The start site of the transcription G is shown as position 1. A gray box indicates an intragenic control region. The pink color highlights TATA-box elements in Hs and Sp U6 promoters. The Proximal Sequence Elements are highlighted in dark-gray. The Distal Sequence Elements (Hs U6: Octamer Motif; Sp U6.1: E-Box) are shown in black. Predicted by using Promo v 3.0.2 (Farré et al., 2003) transcription factor USF1 and USF2 binding sites in LOC119742214 are boxed and highlighted in black. Violet coloring in LOC119742214 indicates the beginning and end of the sequence used as a template of 429 bp to generate sgRNA expression vector. Multiple sequence alignment was constructed using the Clustal W in Mega 11.0.10. Regulatory motifs of Hs U6 and Sp U6.1 were characterized in detail by (Kunkel and Pederson, 1988) and (Sakallah et al., 1994), correspondent. (B) The schematic representation of sgRNA expression vector driven by *Pm*U6 regulatory elements. A guanine (G) nucleotide corresponds to the first nucleotide of the native U6 snRNA. TTTTTT corresponds to the termination sequence.

There are limited numbers of U6 gene studies along echinoderm restricted to the sea urchin (Sakallah et al., 1994). Nevertheless it has been demonstrated in vitro that suUSF (the sea urchin homolog of USF) binds to the E-box sequence (distal sequence elements) and is required for U6 transcription by RNA polymerase III (Li et al., 1994). Using the TFBS prediction tool (PROMO v3, (Farré et al., 2003) we identified potential binding sites for USF1 and USF2 in the LOC109742214 sequence, and determined a 429 bp region upstream of U6 snRNA coding region as a reference sequence to build sgRNA expression vector. The *Pm*U6 regulatory sequence was amplified from genomic DNA merged with *Dll*sg2 followed by polyT-termination sequence (Figure 4B).

*Pm*U6:*Dll*sg2 plasmid vector was coinjected with Cas9 mRNA followed by genotyping analysis to test vector efficiency in vivo (Table S5). The observed plasmid-driven sgRNA the indel frequency was 44.5 ± 2.0 %. This is lower compared with Cas9 mRNA/in vitro transcribed sg2 approach (61 ± 13 %) but is expected based on mosaic expression of plasmid-driven sgRNA.

#### Inducible CRISPR editing of *Dll* loci

Finally we engineered the plasmid expression system to be responsive to DOX and enable inducible in vivo genetic manipulation of *PmDll* loci from a transgene. We built and tested TRE3Gs-regulated Cas9 plasmid vector activated by *Pm*otxG:Tet-ON 3G upon DOX addition, and guided by sgRNA under the control of the endogenous U6 snRNA regulatory elements U6:*Dll*sg2 (Figure 2D, Figure 5).

**Fig. 5.**
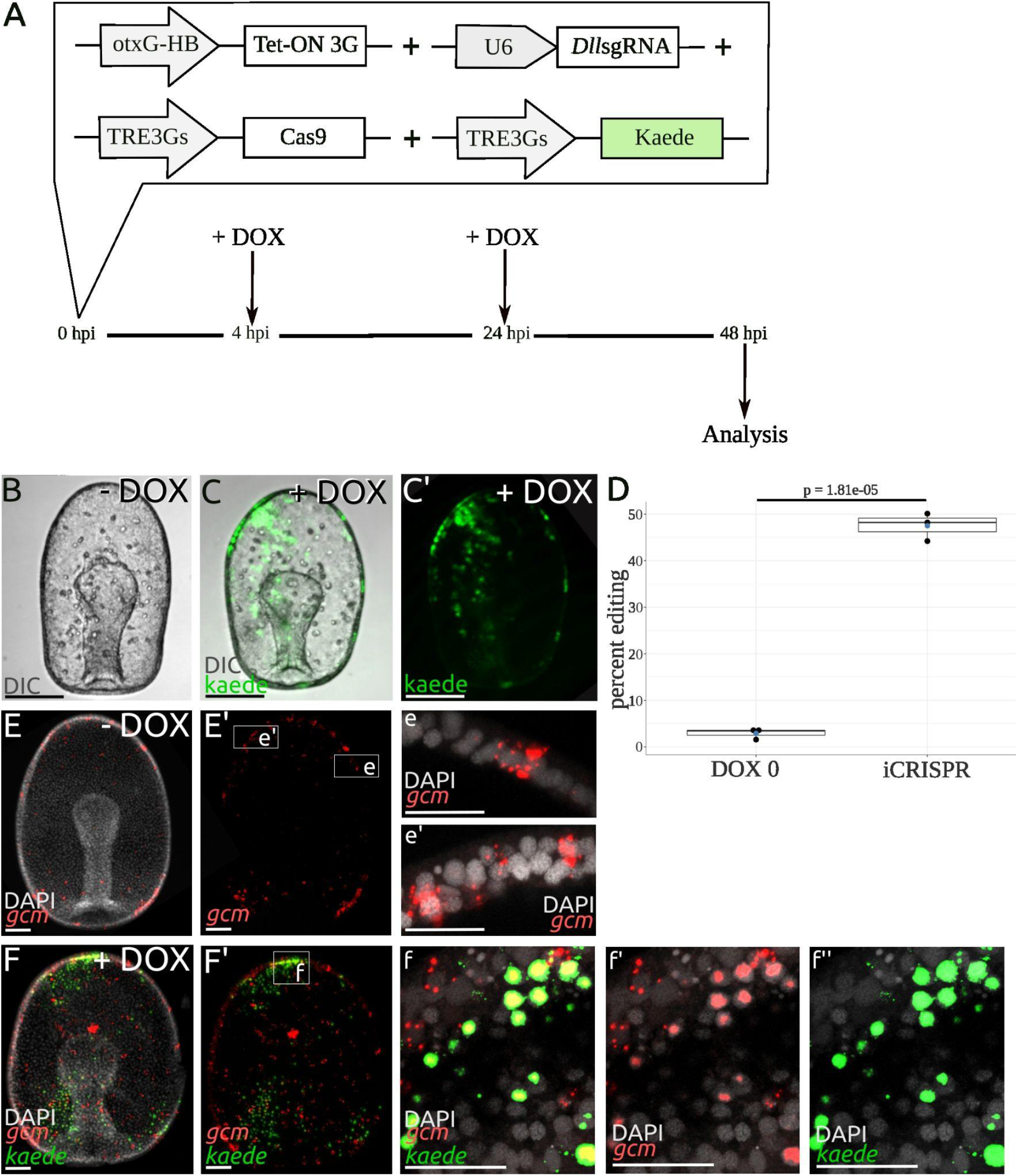
Spatio-temporal control of CRISPR/Cas9 mediated gene editing. (A) Schematic of experimental design. Cocktail of four individual constructs either linearized plasmids or pcr amplicons (Table S5) was injected into the fertilized eggs. OtxG:Tet-ON 3G, TRE3Gs:Cas9 and U6:*Dll*sg2 are iCRISPR constructs. TRE3Gs:Kaede was used as an inducibility reporter - control upon DOX addition. Four hours after injection DOX was added in a dish with injected zygotes with the following drug change at 20 hours later (C-C’, F-f”). Control group (B, E-e’) with injected zygotes received only drug solvent (1/1000 dilution of ethanol) and therefore does not exhibit reporter expression or inducible CRISPR. Induced embryos at gastrula stage were sorted manually based on Kaede expression and subjected to fluorescence WMISH and genotyping analysis. (C-C’) Reporter (Kaede) expression in otxG positive cell populations at 44 hours post induction. Z-projections of confocal stacks of in vivo images. Representative images of control (DOX0) embryo (B) and induced reporter - control along with iCRISPR (C-C’). (D) Indel frequency assessed by single embryo genotypic analysis at the *Dll* loci in transgenic embryos with (iCRISPR) or without DOX (DOX0). The significance of perturbation is calculated by Student’s *t*-test. At least five uninduced and ten DOX+ embryos of three independent biological repeats were genotyped (Table S5). The editing efficiency is calculated using TIDE software (Brinkman et al., 2014). Boxplots showing individual data points (black), mean values (blue) and pairwise P values. (E-f”) Double fluorescence in situ hybridization against *gcm* (red) and *kaede* (green) transcripts in transgenic embryos with (F-f”) or without (E-e’) DOX. (e-e’, f-f”) The expanded view of the region boxed in E’, F’. Z-projections of confocal stacks of images. Scale bars in B-E’: 50 μm; in d-e”: 25 μm.

Considering that the separate transgenic vectors concatenated upon zygote microinjection and inherited as the whole unit into the cell clonality (Hough-Evans et al., 1988; McMahon et al., 1984; McMahon et al., 1985), we injected four individual transgenes either in the linearized plasmids or in the PCR amplicon forms (Table S5). Those are: TRE3Gs:Cas9 linearized (CRISPR part: Cas9), TRE3Gs:Kaede PCR product (inducibility reporter-control), otxG:Tet-ON 3G linearized (spacial control), U6:*Dll*sg2 (CRISPR part: guide RNA) PCR product along with *P. miniata* digested genomic carrier DNA.

Reasoning that the *Dll* zygotic transcription started at 16 hours post fertilization (hpf), we decided to maximize phenotype outcome by inducing Cas9 expression at 4 hours post injection and continue up to 24 hpf (Figure 5A). As expected, no embryos showed Kaede+ expression in the DOX-controls (Figure 5B). Next, using reporter expression as an inducibility marker, Kaede+ embryos (Figure 5 C-C’) were sorted and subjected for genotyping and phenotyping analysis (Figure 5 D, E-f”). 65 ± 13 % of assayed green fluorescent embryos show an average of 47.5 ± 3.0 % mutagenesis in *Dll* loci (Figure 5D, Figure S4 B, Table S5). In our system the spatial induction of mutagenesis is dictated by whatever drives Tet-ON 3G, and thus Cas9 is expressed in mosaic fashion and will be coexpressed with Kaede. Hence we performed fluorescence in situ hybridization to visualize *gcm* expression in regions of *kaede*+ cells (Figure 5 F-f”). A positive assay of a *Dll* CRISPR phenotype will be the expression of adjacent *gcm*+ cells in these regions. At least 25 of uninduced (-DOX) and induced embryos (+ DOX) of three independent biological repeats were subjected to fluorescence double in situ hybridisation. Figure 5 E-F shows representative images of one biological replica. *Gcm* is overexpressed in patches of *kaede*+ clones (compared 5E to 5F) and also significantly expressed in adjacent cells, which is a clear marker of *Dll*-phenotype.

Taken together, our data show that doxycycline-dependent Cas9 induction in vivo (Figure 2D) induces target gene alterations that can recapitulate the effect of the most effective traditional in vitro complexed RNPs knockout approaches (Figure 2A, (Oulhen et al., 2022)). Hence strategies that restrict Tet-ON 3G (rtTA) (Figure 1A) to defined tissue provide a means to produce tissue/cell specific gene disruption. Moreover, using U6-driven sgRNAs targeting several genes will allow manipulation of several genes simultaneously in a given tissue/cell.

## Discussion

The ability to control the timing of transgene expression in a specific tissue is a powerful tool in the study of developmental, regeneration, and cell biology. Such technologies open avenues to explore a myriad of fundamental questions in biology and extend the capabilities of model organisms. Echinoderms present a variety of fascinating traits that require this more fine tuned approach to interrogate the genetic basis of these processes. The model species chosen here, *Patiria miniata*, for example are able to regenerate and have complex developmental and morphogenetic processes that occur post embryonically that are well documented but with little understanding of their genetic and cellular underpinnings. Early embryos of these species are exceptionally experimentally tractable as they are easily injected with antisense reagents, DNA constructs and a range of other molecules and dyes. They are readily cultured and imaged. We sought to extend the capabilities of this species beyond exploration of early developmental processes by designing an inducible gene expression and gene editing tool for use in *P. miniata*.

In order to achieve control of the induction of transgene expression in *P. miniata* we adapted the Tet-ON system. In our constructs we used the commercially available, 3rd generation modified components of the Tet-ON system (Tet-ON 3G and TRE3Gs) to create an spatio-temporal controllable transgenic expression system in the sea star. The spatial control is achieved by utilizing cell/tissue specific regulatory elements, whose repertoire is expanding due to available ATAC-seq data during developmental and regeneration time courses. The transgene expression itself is placed under chemical control.

The Tet ON/OFF system, has been developed over 25 years from components of the bacterial transposon Tn10-encoded Tet operon (Hillen and Berens, 1994) and has been successfully applied in numerous tissue culture systems as well as in a broad range of species (yeasts, protozoans, insects, amphibians, plants and mammals). However, there are limited examples of versions adopted for use in echinoderms or to our knowledge in other marine organisms.

In this paper as a proof-of-concept demonstration we use spatio-temporal control of *Sp*Cas9 endonuclease expression to create the inducible CRISPR - mediated gene editing system in *P. miniata*. We showed that the Tet-ON system effectively induces gene expression in the sea star, *P. miniata* at 15°C and in sea water. DOX induction at either 5 or 10 μg/ml effectively induces expression of our reporter gene under the control of TRE3GS when its transcription activator (Tet-ON 3G) is under the regulatory control of the *P. miniata* enhancer, otxG. Our control experiments confirm that there is no expression leakage in the absence of DOX induction. Expression is driven clonally, as expected, since plasmid based reporters are known to incorporate into cell lineages only after one to several cell divisions. The reporter expression driven by the two component plasmid system once induced by DOX was spatially equivalent to that of a single plasmid with otxG enhancer. The transgene transcripts are expressed in the otxG+ in predominantly ectoderm and in the endomesoderm from early blastula stage continues to gastrula. The applications of this two plasmid Tet-ON system are in principle tunable to a vast range of ectopic gene expressions, such as (fluorescence reporter, calcium indicators, recombinases, endonucleases), targeted cell ablation (Labbaf et al., 2022), controllable functional inactivation of genes by dominant negative mutants. These tools could accelerate the use of sea star as a scalable invertebrate model in developmental, regeneration, aging studies and cancer biology.

Previous studies used CRISPR targeted genome editing in echinoderms to alter either structural (Lin et al., 2019; Mellott et al., 2017; Wessel et al., 2020) or regulatory (Pieplow et al., 2021) genes, and to create the first stable transgenic sea urchin lines (Liu et al., 2019; Vyas et al., 2022; Yaguchi and Yaguchi, 2022). However, these were not tunable for in vivo spatio-temporal regulation.

We first showed that CRISPR/*Sp*Cas9 gene editing works efficiently when either RNPs or Cas9 mRNA with guide RNAs were injected into the zygote by targeting the well characterized *Delta* gene. With sgRNA targets to exon four shared by both splice variants we observed the well known phenotype of the overexpression of the transcription factor *gcm*. In normal embryos, *gcm* is expressed at low levels in a “salt and pepper pattern” of a *gcm+* cell surrounded by *gcm-*cells, particularly evident in the anterior ectoderm. We have previously reasoned that this is due to a lateral inhibition mechanism. When early embryos are injected with antisense morpholinos blocking the translation of *Delta* (Fig. S2), *gcm* is expressed at higher levels and in adjacent cells. Either method of introducing CRISPR components (RNP or RNAs) phenocopies this *gcm* misexpression. We additionally quantified the percentage of gene edits in the *Delta* genomic locus at the single embryo resolution using Tracking of Indels by DEcomposition (TIDE) software. This showed, in case of RNPs, the mutation efficiency is 68.3 ± 26.5 %, while the transcribed Cas9 and guides produced 61 ± 13 % editing. This is in keeping with other work in echinoderms, for example editing efficiency from sea urchin *Alx1* promoter mutagenesis experiments using Cas9 mRNA with “home-made” guide RNAs were 53 and 69.2% (Pieplow et al., 2021); in case of RNPs targeting sea urchin *FoxY*, the mutation rate was 71% (Oulhen et al., 2022).

In order to couple this system to the two plasmid component Tet-ON system we needed to introduce both Cas9 and guide RNAs on expression constructs. We reasoned that the guide RNA could be ubiquitously expressed and the Cas9 expressed under spatiotemporal control. We thus designed a plasmid expression vector using the *P. miniata* snU6 upstream non coding regulatory region which uses pol III promoter optimal for driving expression of short RNAs. At this point we do not investigate the cell type expressing snU6 but reasoning that as snU6 genes are highly conserved we assume that they are active in all cell types like in mammalian species (Sandy et al., 2005). We then introduced Cas9 on TRE3GS DOX inducible construct which places it under the regulatory control of DOX and rtTA (Tet-ON 3G). Using the same concentrations of DOX we induce the expression of Cas9 from this construct along with the reporter to mark the clones of *Cas9+* cells. We observe a similar spatial localization of Kaede in this experiment as the initial Tet-ON experiment (Figure 1D-E’ and Figure 5C-C’). This is expected as the experiment is identical except now with the addition of the TRE3GS:Cas9 and U6:*Dll*sg plasmids. We show that *gcm* is indeed expressed in adjacent cells only within the Kaede+ clones confirming that *Delta* expression is disrupted. The percentage of mutagenesis at the expected *Dll* loci in these embryos was on average of 47.5 ± 3.0 %. This is lower than when RNAs are injected into the zygote which is probably due to the mosaic incorporation. However, continuously expressing several guides targeting the same gene and rtTA (Tet-ON 3G) from the strong transcription regulatory elements may provide higher levels of editing in the transgenic cells.

We show that we can now achieve spatial and temporal control of gene editing, which opens up a wealth of opportunities to explore post embryonic development, regeneration and cell biological processes in this species. The approach we show here can be adapted for use in other species of echinoderms and will also extend experimental possibilities tremendously. There remain some limitations. The first is that any editing will be achieved in clones of cells. This will limit the ability to identify gross phenotypes. Coinjecting a reporter construct or designing another coexpression system will ensure that the clones of Cas9 expressing cells are identified and traceable. The cells can then be subjected to more fine resolution assays, such as changes in gene expression, protein localization using immuno-histochemistry and/or live imaging of cell behaviors. Another significant limitation is that the extent of gene editing in the individual clones and hence the penetrance of the phenotype is not known. We do however show a high percentage of mutagenesis in these clones, suggesting the driving expression of Cas9 and sgRNA is a very effective approach to achieve editing. These experiments should however be coupled with controls that assess perturbation of the target gene. Moreover, future dissections of *Pm*U6 regulatory elements and establishment of a minimal promoter would allow for small RNA expression, such as siRNAs and shRNAs, under DOX control.

A future direction is to establish lines of these mosaics to generate F1 and F2s and thereby to establish stable lines where the TRE3Gs:Cas9 transgenes are present in all cells. Cell/tissue specific enhancers control production of Tet-ON 3G and express Cas9 only when DOX is added. The improved reproducibility of experiments will propel the utility of this species for understanding a diversity of biological processes.

## Materials and Methods

### Embryo cultures

Adult *Patiria miniata* were obtained from California, US (Pete Halmay or Marinus Scientific) and were used to initiate embryo culture following the established protocol (Cheatle Jarvela and Hinman, 2014). Embryos and larvae were cultured in artificial sea water and fed algal cultures of *Rhodomonas lens* (AlgaGen, LLC).

### Plasmid constructs

Plasmids (otxG:Tet-ON 3G; TRE3Gs:Cas9; TRE3Gs:Kaede; U6:sg2; Table S1) were built by Gibson cloning (NEB, Cat# E2611) following standard protocols. In-house expression vectors were used as a backbone, and as well as a source for otxG, HB and SV40 polyA. TRE3Gs and Tet-ON 3G were amplified by PCR from AAVpro Tet-One Luc Control Vector (Clontech, Cat# 634311), Kaede from in-house recombineering cassette (Zheng et al., 2022), hCas9 from addgene Cat# 72602; U6 from *P. miniata* genomic DNA, sg2 from DNA template for IVT sgRNA reaction. To obtain a linear DNA template plasmid TRE3Gs:Cas9 was linearized with SapI and otxG:Tet-ON 3G with SalI.

### RNA in vitro transcription (IVT)

### IVT capped mRNAs

5’ - capped and 3’- polyadenylated hCas9-2xNLS mRNA was transcribed in vitro using mMessageMachine T7 (ThermoFisher, Cat# AM1344) and Poly(A) Tailing (Applied Biosystems, Cat#AM1350) kits. Unincorporated nucleotides were removed by precipitation with LiCl.

### IVT sgRNAs

DNA template for IVT reaction was generated by overlap extension pcr using Phusion-HF DNA Taq polymerase. Two assembly oligos (IDT, ultramer DNA oligos) were used in reaction: one oligo contains target specific sequence - overlap, other oligo contains overlap - Cas9 Scaffold (Table S4). The sgRNAs were synthesized by T7 RNA polymerase using the MegaShortScript T7 transcription kit (ThermoFisher, Cat# AM1354) following the manufacturer’s instructions. Then the sgRNAs were purified using the MEGAclear kit (ThermoFisher, Cat# AM1908) with ammonium acetate ethanol precipitation.

### Live imaging

Before imaging the embryos were immobilized in 500 mM high-salt sea water for 1–2 min to remove the cilia, then mounted on the slides. Imaging was performed by using the Andor Revolution XD spinning Disk confocal microscope with Andor IQ3 system using cDIC and c488 channels.

### Whole mount colorimetric in situ hybridization (WMISH)

WMISH using digoxigenin-labeled antisense RNA probes followed previously published protocols (Hinman et al., 2003; Yankura et al., 2010). Briefly, embryos were fixed in 4% paraformaldehyde in MOPS buffer (0.1M MOPS pH 7.5, 2mM MgSO4, 1mM EGTA, and 800mM NaCl) overnight at +4℃, then transferred to 70% ethanol solution. Hybridization of *gcm* digoxigenin-labeled antisense riboprobe (final concentration of 0.2 ng/ml) was performed at 58 °C for 5 days. Detection was performed using an anti-digoxigenin AP-conjugate antibody (Roche, Cat# 11093274910) followed by an NBT/BCIP reaction (Roche). Images of whole-mount samples were taken using a Leica DMI 4000B microscope equipped with a Leica DFC 420C camera. At least fifty samples per two independent biological replicas were examined.

### Whole mount double fluorescent in situ hybridization

Double FISH experiments were performed as previously described (McCauley et al., 2013) using digoxigenin-labeled antisense *kaede* riboprobe and dinitrophenol-labeled antisense *gcm* riboprobe (Table S1), which were hybridized simultaneously. The probes were detected using anti-DIG POD-conjugated antibody (Roche, Cat# 11207733910) and anti-DNP HRP-conjugated antibody (Perkin Elmer, Cat# FP 1129) followed by tyramide signal amplification (Akoya Biosciences: TSA Plus Fluorescein Cat# NEL741E001KT, TSA Plus Cyanine 3 Cat# NEL744001KT). The labeled embryos (DAPI, Fluorescein, Cy3) were imaged using a Zeiss LSM 880 scanning laser confocal microscope (405, 488, 561 channels). At least ten samples per three independent biological replicas were examined.

### RNA extraction and QPCR

RNA from 25-50 embryos was isolated with the GeneElite (RNA miniprep kit, Sigma Cat# RTN350) kit with DNAse I treatment (ThermoFisher, Cat# EM0521) on column followed by additional DNAse treatment using DNA-free DNA Removal kit (Invitrogen, Cat#AM1906). cDNA synthesis was performed using the Verso cDNA Synthesis kit (ThermoFisher, cat# AB1453). qPCR was performed with home-made SYBR green I (Invitrogen, Cat# S7563), ROX Reference Dye (Invitrogen, Cat# 12223012), and Phusion High-Fidelity Taq polymerase (ThermoFisher, Cat# F530) master mix using QuantStudio3 (ThermoFisher) real-time PCR systems. Efficiency of *kaede* primers was calculated using standard curves in duplicate with four serial dilutions (Table S3). Transcripts were normalized to *laminin2b receptor* (Cary et al., 2020). Three biological repeats and three technical replicates were performed (Table S3). The relative fold of *kaede* expression was calculated using the delta Ct method. Graphs and statistical analysis (Student’s t-test) were performed using RStudio.

### Genotyping

The genomic DNA of individual control or *Dll*-KO embryos (sorted manually based on injection marker or on reporter expression) were extracted with 20 μl of Quick Extract ™ DNA Extraction Solution (Lucigen, Cat# QE09050) according to the manufacturer’s instructions. Four microliters of the extraction mix was then subjected to PCR amplification of the targeted genomic DNA region using recombinant Taq DNA polymerase, (ThermoFisher, Cat# EP0404). Then PCR products were column purified (Zymo Research, Cat# D4004) and Sanger sequenced (Table S1, Table S4, Table S5). Spectral data (.ab files) of these sequences were subjected to TIDE analysis (Brinkman et al., 2014) with default parameters. Edited chromatograms were aligned against unedited control sequences for indel decomposition analysis. To exclude false positive results due to the errors based on PCR reaction, Sanger sequencing the control sequences representing Cas9 only or DOX minus (DOX0) were subjected to TIDE analysis against control sequences. At least two independent biological repeats were performed for all knockout experiments. The percent editing in *Dll*-KO embryos was quantified relative to unedited embryos. Graphs and statistical analysis (Student’s t-test) were performed using RStudio.

### Microinjection

All perturbing reagents used in this study were delivered into the fertilized eggs by microinjection following established protocols (Cheatle Jarvela and Hinman, 2014). Healthy morphant embryos were subsequently isolated for quantitative and qualitative studies. hCas9 mRNA was delivered at concentration of 1μg/μl of injection solution; home-made sgRNAs at 250 - 500 ng/μl; RNPs complexes at 1 to 5 molar ratio of *Sp*Cas9 2xNLS recombinant protein (Synthego) and chemically synthesized sgRNAs with protecting groups (Synthego) at concentrations of 1.2μM Cas/ 6μM sg; 1.9μM Cas/ 9.3μM sg; 3.75μM Cas/ 18.6μM sg. As *P. miniata* fertilized eggs are sensitive to glycerol, RNPs complexes were subjected to dialysis in 120mM KCl solution before injection. The control injections of hCas9 mRNA only, sgRNAs only, and *Sp*Cas9 nuclease only were delivered at the same concentrations as for experimental design. The total amount of injected DNA either as circular, linearized, or as a PCR product was 10-15 ng/μl.

### Drug treatment

Doxycycline hyclate (Sigma, Cat# D9891) was dissolved in 100% ethanol at 10 mg/ml for storage and maintained as a stock solution in the dark, at α20 °C. Embryos at a density up to 30 per four ml of sea water in small petri dishes were treated with 5 and 10 μg/ml DOX for 24 - 48 hours. Sea water and DOX were changed every 24 h to avoid break-down of drug. Due to light sensitivity of the drug, embryos were protected from light during DOX treatment.

### Software

CRISPR guide RNAs were designed using CHOPCHOP v3 (Labun et al., 2019). Fiji image software (Schindelin et al., 2012) was used for preparing Z-projections of stacks of confocal optical sections. Figures were constructed using Inkscape 1.2.1 and GIMP 2.10.32. Graphs and statistical analysis were performed using RStudio. Mutation efficiency was calculated using TIDE (Brinkman et al., 2014). Sequence chromatograms were analyzed using FinchTV. Multiple sequence alignment was performed using Mega 11.0.10 (Kumar et al., 2018) and visualized by Jalview (Waterhouse et al., 2009). Putative transcription factor binding sites in DNA sequence were identified using PROMO v3 (Farré et al., 2003) based on TRANSFAC v8.3 weight matrices database.

## Acknowledgements

We thank all authors of two inspirational papers from Rosenthal (Crawford et al., 2020) and Gladyshev (Nguyen and Gladyshev, 2020) labs without which this work would be impossible. We would also like to thank John Lee Andrade for help with ATAC-seq data, Jessalyn Grant-Bier for help with troubleshooting tips and Cheryl Telmer for helpful feedback during the preparation of the manuscript.

## Supplementary Data

**Table S1.**
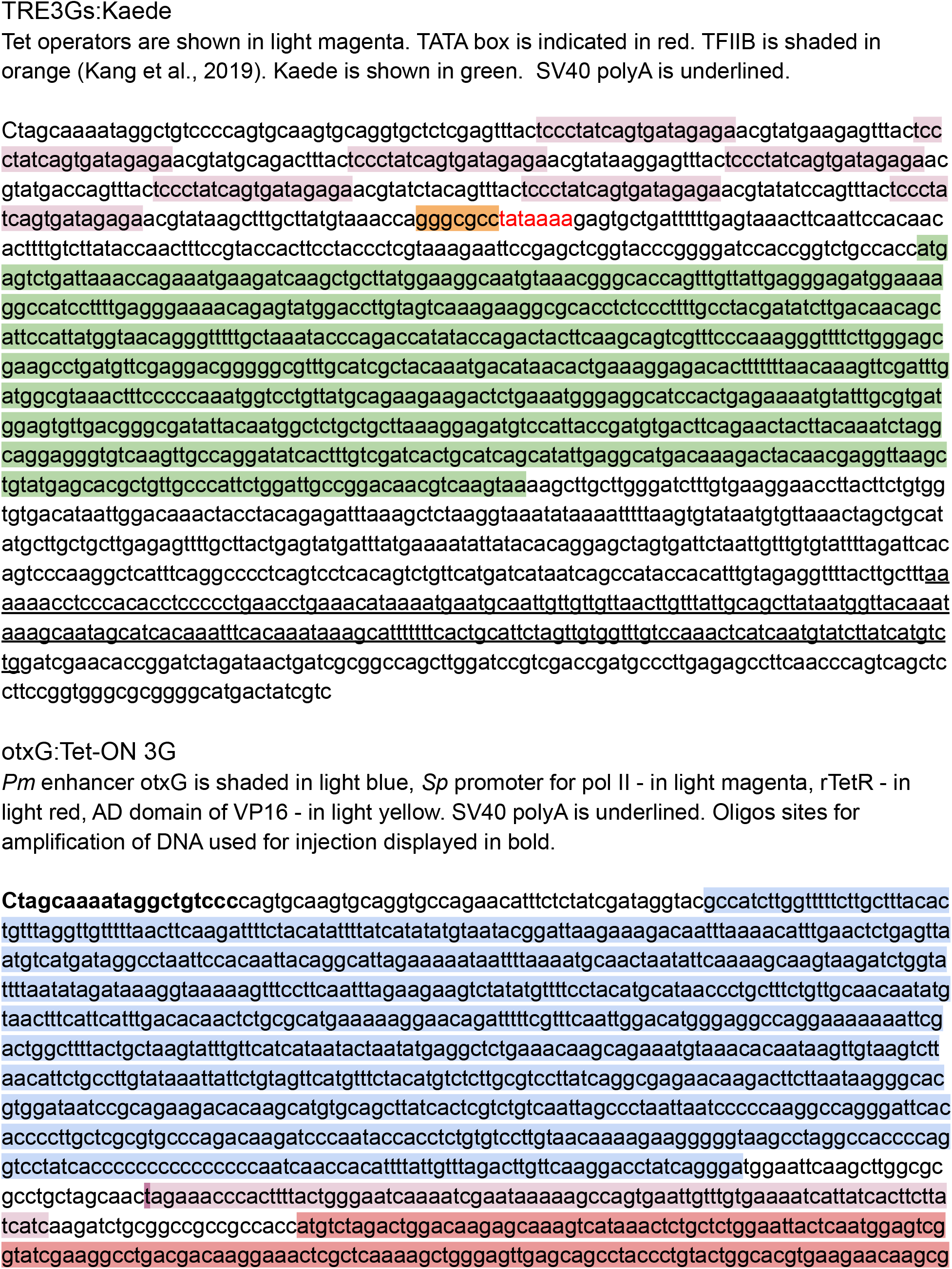
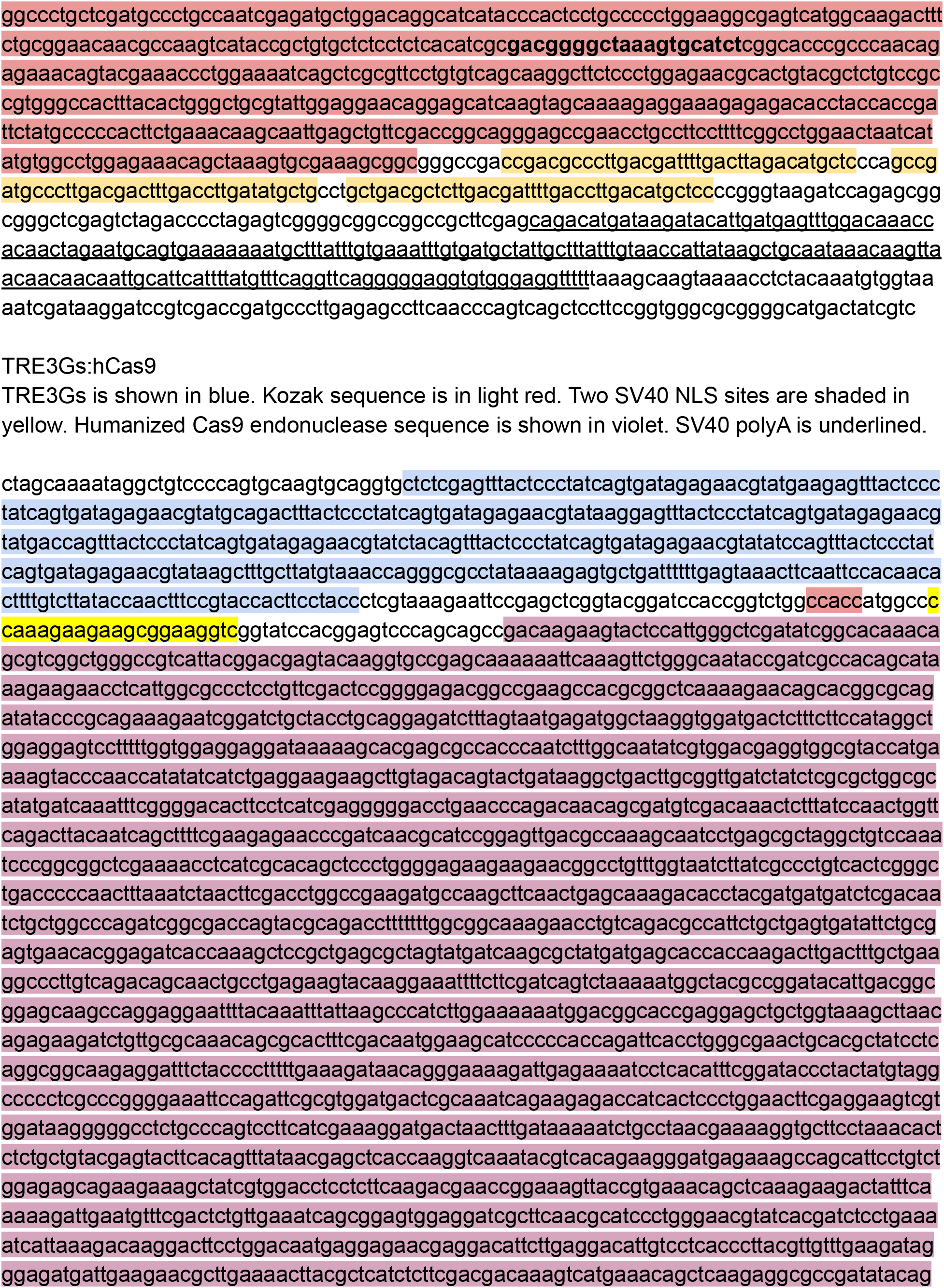
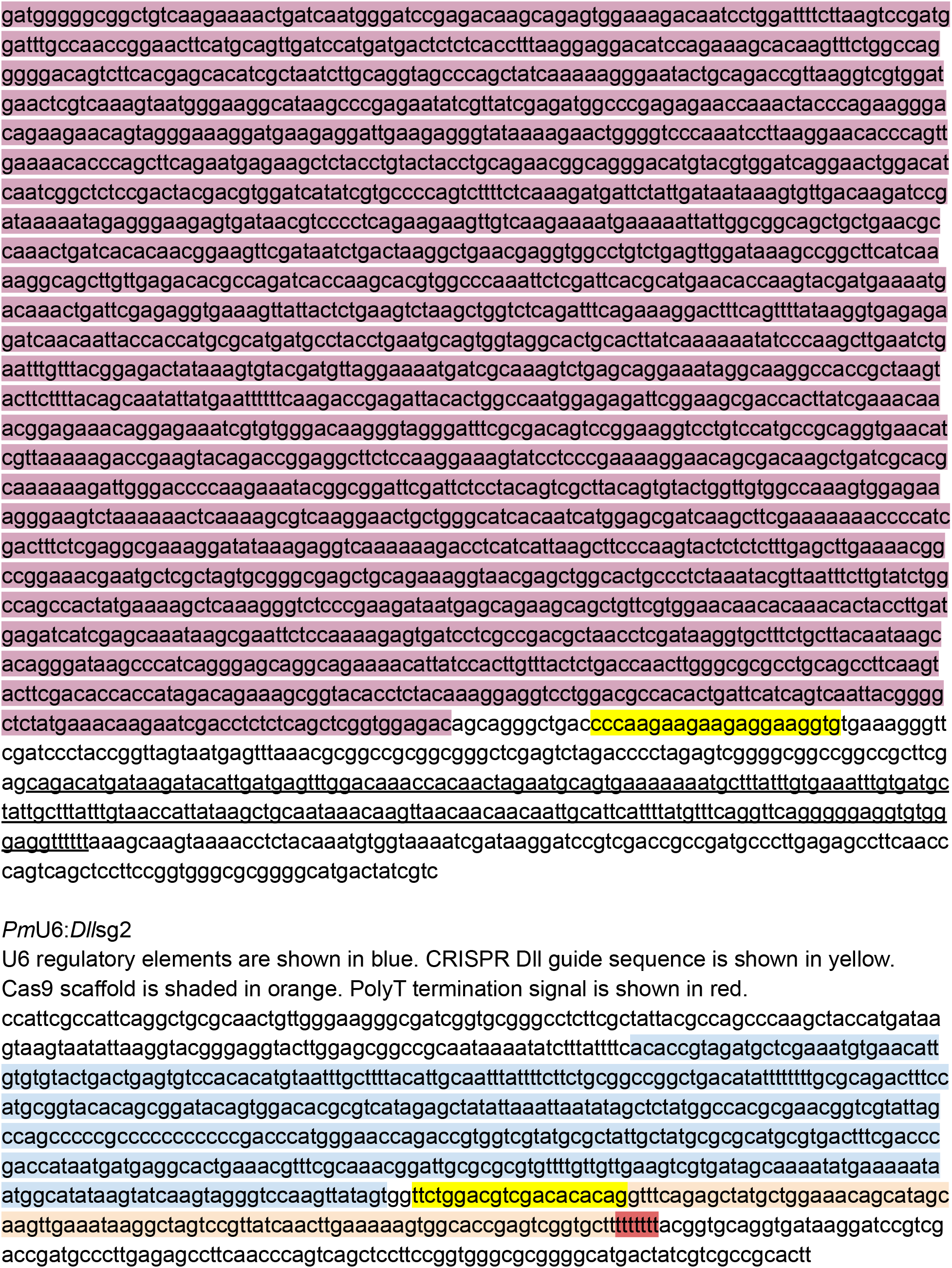
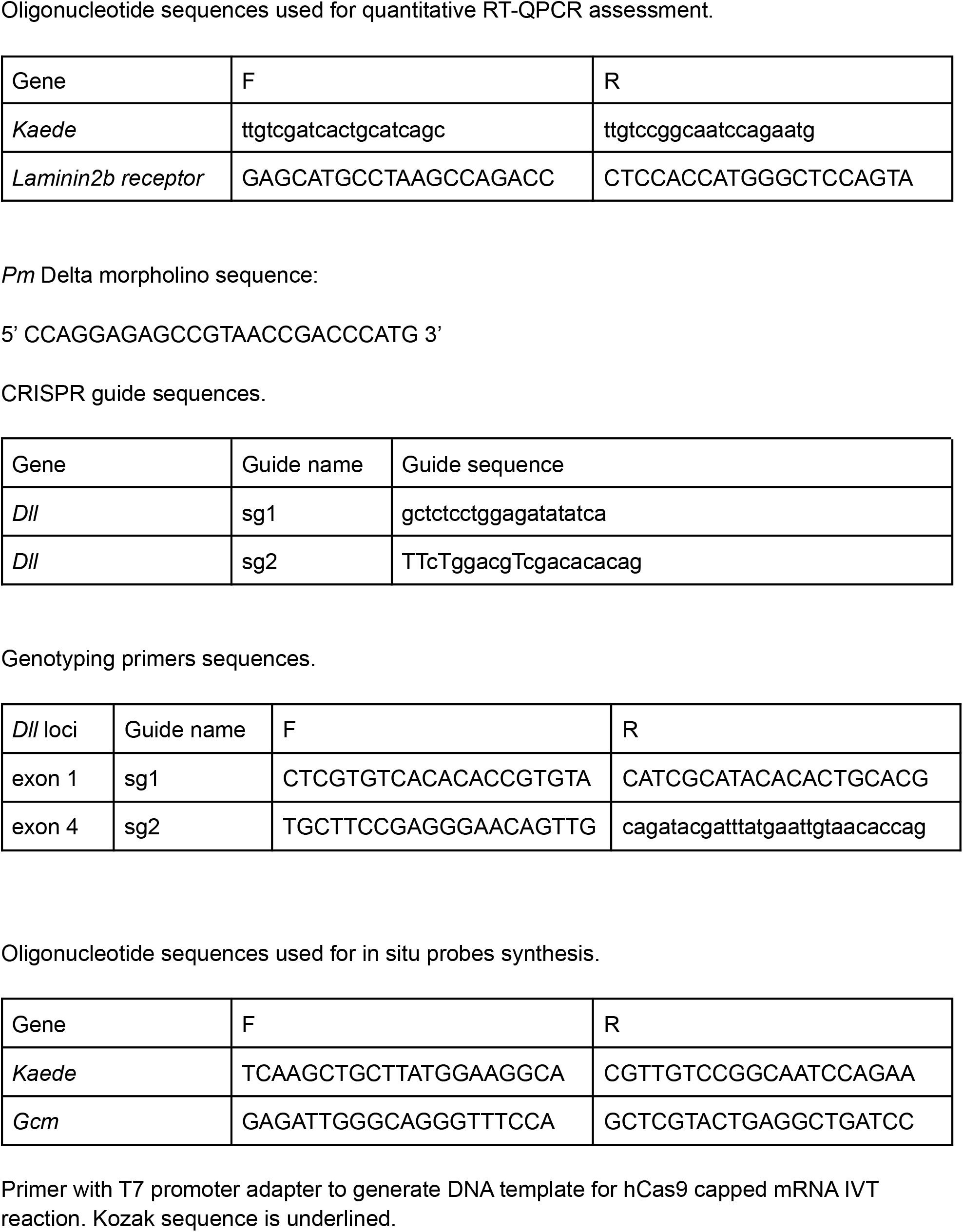
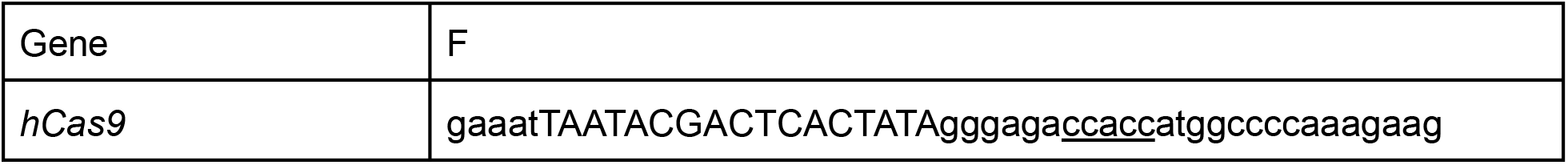
Annotated sequences of used vectors.

**Table S2**. Reporter assays

**Table S3**. QPCR data

**Table S4**. Genotyping RNPs & Cas9 mRNA/sg2

**Table S5**. Genotyping Cas9 mRNA/U6:sg2 & iCRISPR

**Fig. S1.**
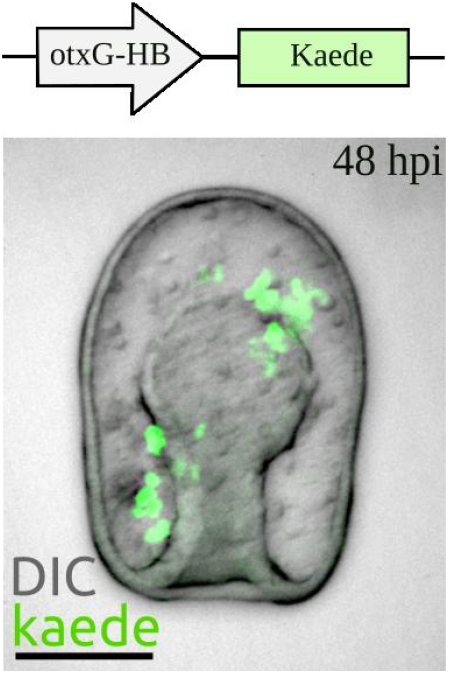
Kaede expression driven by *Pm*-otxG enhancer. The reporter is expressed clonally and fated predominantly ectodermal and the endomesodermal cells. OtxG was previously identified as the cis-regulatory element responsible for control of *Am*otxβ1/2 expression during blastula and early gastrula stage (Hinman et al., 2007). Expression construct contains the RNA polymerase II basal promoter for the sea urchin hatching enzyme (HB, Wei et al., 1995) and a Kaede reporter. Representative image of embryo showing Kaede+ ectodermal cells under *Pm*-otxG enhancer. Z - projection of confocal stack of in vivo image. Scale bars: 50 μm.

**Fig. S2.**
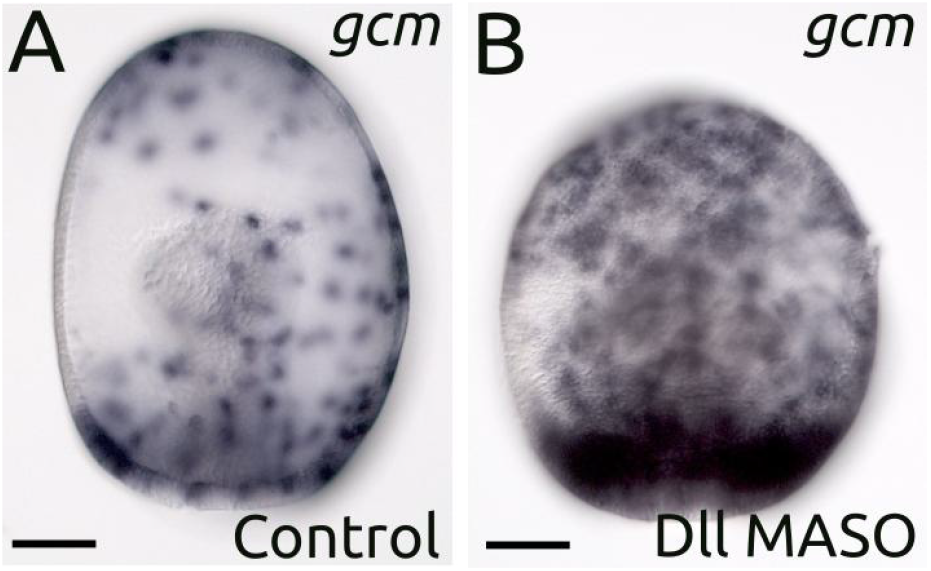
Knockdown of Delta (*Dll*) with an antisense morpholino (sequence information in Table S1) results in massive ectodermal *gcm (glial cells missing* transcription factor) overexpression compared to control morpholino injection. Images are colorimetric WMISH with the probe to the *gcm* at gastrula stage. (A) Control embryo showing “salt and pepper” *gcm* expression. (B) *Dll* MASO morphant. Data are representative of two biologically independent replicates consisting of at least ten embryos each. Zygotes were injected with MASOs (GeneTools) following the protocol by Cheatle Jarvela and Hinman ((Cheatle Jarvela and Hinman, 2014). The GeneTools standard control MASO was used as a control morpholino injection. Scale bars: 50 μm.

**Fig. S3.**
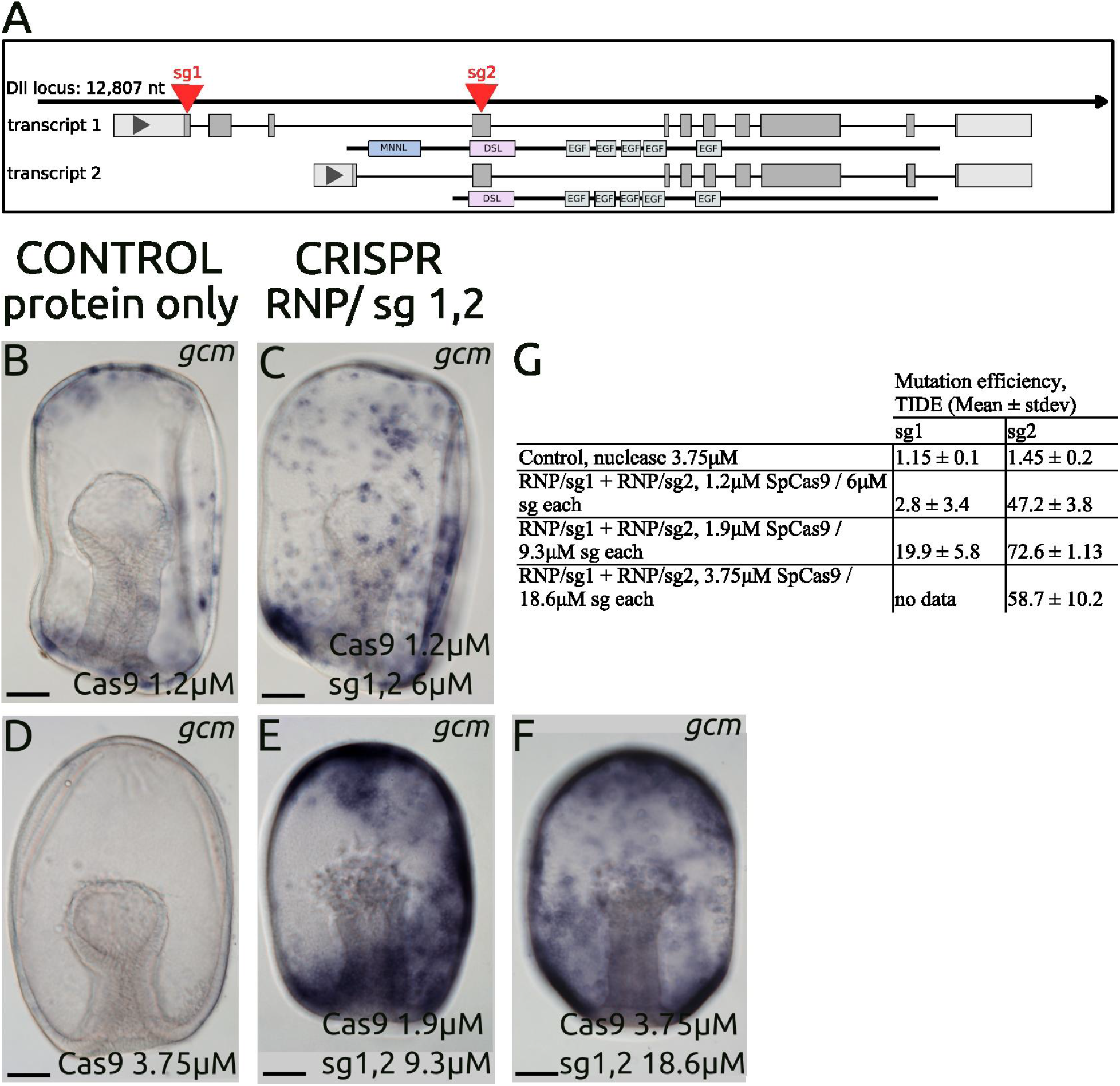
Dose-dependent RNP editing. (A) Schematic of Delta loci and Cas9 targeting sites (sg1, sg2). (B, C) RNP of 1.2 μM Cas9 protein (B, protein only) complexed with 6 μM Synthego synthesized guide RNAs each (C). (E) RNP of 1.9 μM nuclease complexed with 9.3 μM sg1, sg2 each. (F) RNP of 3.75 μM nuclease (D, protein only as a control) complexed with 18.6 μM sg1, sg2 each. Phenotypic analysis of control (B, D) and *Dll*-KO (C, E, F) embryos visualized by WMISH against *gcm*. (G) Table of mutation efficiencies calculated by TIDE. Data are presented as mean values ± standard deviation. At least four individual embryos of one biological repeat were genotyped for each experiment. RNP of 1.9 μM nuclease with 9.3 μM sg2 was subjected to the following analysis. Scale bars: 50 μm.

**Fig. S4.**
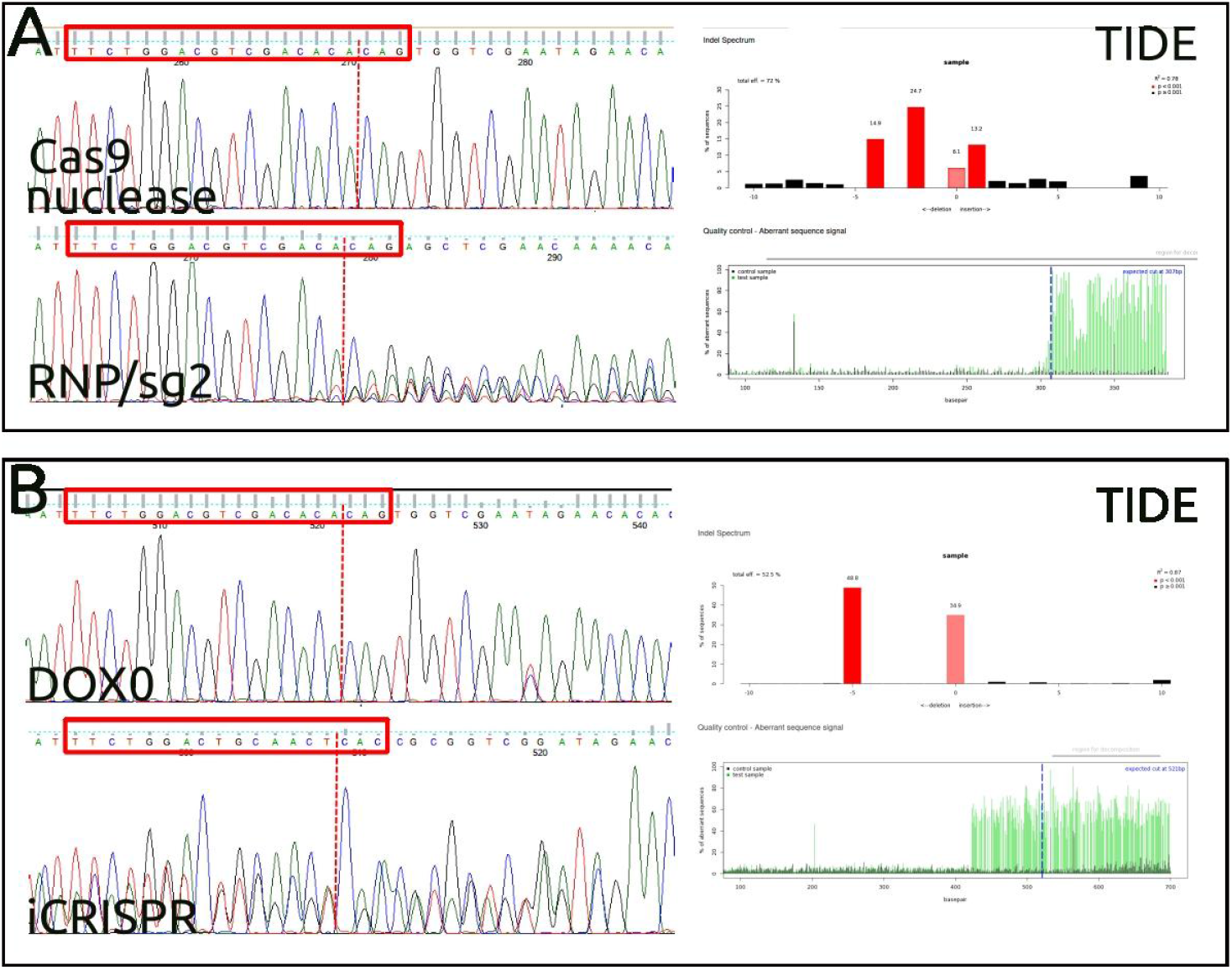
Genotyping of *Dll*-KO embryos. Sequence chromatograms of PCR products of unediting (Cas9 nuclease, DOX0) and editing (RNP/sg2, iCRISPR) cell populations at single embryo resolution. Red dotted line indicates the position of the Cas9 canonical cut site. Guide sequence is indicated in the red box. TIDE (Tracking of Indels by DEcomposition) analysis showing the indel spectra and aberrant sequences of editing versus unedited cell populations. Upper panel shows the indel frequencies within ± 10 bp from the theoretical gRNA cleavage site. Lower panel depicted PCR sequences aberrations; the theoretical cut site is indicated by a blue line. (A) Editing at *Dll* (exon 4) loci by RNP complex. (B) Induced upon DOX addition editing of *Dll* loci (exon 4) in otxG+ cell populations.

